# Multivesicular release at a small central excitatory synaptic contact

**DOI:** 10.1101/2025.08.06.668911

**Authors:** Boris Barbour, Ann Lohof

**Author notes:** **Background** This report (except this front page) was written in 2001 and has not been changed since. It is based upon experiments performed in 1997--1998. It represents a preliminary (but sufficient) analysis of recordings of cerebellar granule cell--Purkinje cell pairs directly demonstrating multivesicular release at what are likely (and in a few cases shown) to be single excitatory synaptic contacts. **Author contributions:** The project was conceptualised and designed by BB, who also performed the majority (19/20) of the recordings shown, the analysis and wrote the MS. AL contributed to the experiments, including one connected pair recording and reviewed the MS.

## Abstract

We set out to study the properties of excitatory synaptic transmission at a central connection likely to consist of a single small synaptic contact. In the cerebellar cortex, molecular layer interneurones extend sparse thin dendrites in a parasagittal plane. The nonbranching parallel fibres traverse at right angles to the dendritic plane, forming en passant excitatory synapses. This arrangement makes it very likely that any synaptic connection between a parallel fibre and an interneurone will involve only a single contact. We obtained 20 paired recordings of synaptically connected granule cells and interneurones in cerebellar slices of immature rat, recorded at room temperature. Three of these pairs were successfully labelled using biocytin and a single point of contact between them was indentified. In 19/20 recordings, multiple release events corresponding to asynchronously released vesicles were frequently resolved following a single action potential. The frequency of these events could not be accounted for by spontaneous EPSCs, whose frequency was very low. We conclude that at this excitatory synaptic contact an action potential can release more than one vesicle.

---

The principal physical traces of memory and learning are modifications of the properties of transmission at distributed subsets of synapses. Although understanding these modifications at a synaptic level has been the focus of intense effort, these studies have been severely hampered by the paucity of information regarding the basic functioning of central synapses. An example of our lack of knowledge is the question of how many vesicles are released at central synapses per action potential (Auger and Marty, 2000). For many years it was assumed that an action potential at the smallest central synaptic contacts could release no more than a single vesicle of neurotransmitter (Korn et al., 1982). However, some evidence (Tong and Jahr, 1994), particularly for an inhibitory synapse (Auger et al., 1998), indicates that ‘multivesicular’ release can occur. The majority of central synapses are excitatory, but no clear demonstration of multivesicular release exists for these synapses. Indeed, monovesicular release has been established in some special cases (Gulyas et al., 1993; Stevens and Wang, 1995; Silver et al., 1996). Obviously, the release of multiple vesicles is often observed at synaptic connections, but it has proved difficult to rule out that those connections comprise multiple contacts.

As described in the abstract, the parallel fibre-interneurone synapse in the cerebellar molecular layer provides an ideal preparation in which to examine the question of multivesicular release (Palay and Chan-Palay, 1974). These synaptic connections should involve only single contacts and recordings display excellent signal-to-noise ratios. We therefore set out to record from such synapses in paired recordings, to ensure the selective stimulation of the presynaptic granule cell and to allow labelling of both cells. These experiments were made difficult by the low probability of finding such connections in paired recordings. Nevertheless, a small number of pairs were successfully recorded and they show that multivesicular release can occur at this synapse.

## Materials and Methods

### Slice preparation

2--3 week-old rats were killed by decapitation using a small animal guillotine. The cerebellum was quickly dissected out into ice-cold saline with the following composition (concentrations in mM): NaCl, 125; NaHCO_3_ 26; KCl 2.5; NaH_2_PO_4_ 1.25; CaCl_2_ 2; MgCl_2_ 1; D-glucose 25. Transverse slices (350 µm) oriented at a 20° angle to the long axis of the cerebellum were prepared using a DTK-1000 Microslicer (DSK EM Corp, Kyoto, Japan). Recordings were carried out at room temperature in saline of the following composition. For a few experiments (in mM), ascorbic acid, 0.4; myo-inositol, 3; Na-pyruvate, 2 were added to the experimental saline. Experiments were carried out in the presence of a GABA_A_ antagonist (20µM bicuculline methochloride or 10 µM gabazine) and 50 µM D-APV. Slices were viewed through a Zeiss Axioskop (Carl Zeiss France SAS, Le Pecq, France) with a 0.75NA 40x water immersion objective and a 0.9NA condensor. Green illuminating light was selected with a bandpass interference filter (wavelengths 500 ± 30 nm). A video image was obtained with a 2/3” CCD camera and displayed after electronic contrast enhancement by a home-made circuit. Drugs and chemicals were obtained from Sigma-Aldrich Chimie S.a.r.l. (Lyon, France) or Tocris (Bristol, UK)

### Recording and finding synaptic connections

The following protocol was adopted to increase the probability of finding synaptically connected pairs. First an interneurone was whole-cell patch-clamped, using an Axopatch 200A (Axon Instruments, Union City, CA). The pipette solution contained (in mM): K-gluconate, 145; KCl, 5; HEPES, 10, EGTA 0.2 (for granule cells) or 1 (for interneurones); MgCl_2_, 2; Na_2_ATP, 2; biocytin, 13. In some experiments Na_3_GTP, 0.4, Tris_2_phosphocreatine, 10; reduced glutathione, 5 were also added, with the amount of K-gluconate reduced to maintain osmolarity. The series resistance was in the range 10--39 MΩ (mean 26 ± 7.5 MΩ), but series resistance compensation of nominally 60--80% (reported by the potentimoter of the amplifier) was usually applied.

In order to locate viable connected granule cells, chemical screening was carried out, using 100 µM kainate (dissolved in [in mM]: 150, NaCl, KCl, 2.5; HEPES, 10; NaH_2_PO4, 1.25, CaCl_2_, 2, MgCl_2_, 1) pressure ejected from a patch pipette. It was necessary to be able to move the microscope independently of the recorded cell, because the search for granule cells was carried out from a few hundred microns to more than 1 mm distant from the interneurone. The locations where brief puffs of kainate induced immediate EPSC activity in the interneurone marked the positions of the ends of the granule cell dendrites (rather than the cell body). Nearby granule cells were then screened individually using loose cell-attached stimulation (the kainate-containing pipette was changed to a similar one containing the same solution without kainate). Pulses of +1- -2 V and 1--2 ms long were applied, though these parameters would not now be recommended (Barbour and Isope, 2000), since they are unnecessarily damaging. The signal-to-noise ratio of the recording allowed immediate recognition of the connected granule cell. With a clean electrode containing intracellular solution (described above), an attempt would then be made to establish a whole-cell patch-clamp recording of the connected granule cell.

The granule cell was recorded and stimulated using an Axoclamp 2B (Axon Instruments, Union City, CA), switching between voltage-clamp and current-clamp. Neutralisation of the electrode capacitance was applied. The ‘bridge’ was not balanced. The resulting (small) error did not prevent detection of the elicited action potential. Except when being stimulated, granule cells were voltage clamped. In preparation for stimulation, the granule cell was switched from voltage-clamp to current-clamp. An action potential was then elicited by the immediate injection of a depolarising current. Shortly after the action potential, the granule cell was switched back to voltage clamp. This hybrid protocol offered a number of advantages: it prevented unwanted action potentials from disrupting the programmed intervals; it allowed control of the interstimulus membrane potential; it made the stimulation more reproducible, by imposing a uniform starting potential for each stimulation. Granule cells were stimulated at about 0.3Hz to emit pairs of action potentials with various intervals (about 5--65 ms).

Correction has been made for junction potential errors (about 11mV). Holding potentials were between -73 and -81mV for the granule cells and -81 mV for the interneurones. Recordings were filtered with a 4-pole Bessel filter (built-in) set to a corner frequency of 10KHz and sampled at 50KHz. The traces shown in the figures have undergone additional Gaussian filtering (σ = 50 µs).

### Histology

After the experiment, the pipettes were carefully withdrawn in order to form outside-out patches, if possible, and thus retain the biocytin within the recorded cell. The slice was fixed in 4% paraformaldehyde in PBS. The revelation of the biocytin followed standard procedures. After washing the fixative and blocking endogenous peroxidases with 10% methanol/1%H_2_O_2_ in 0.4% triton-X-PBS, the slice was incubated with a streptavidin-HRP complex (Vectastain ABC kit, ABCYS, Paris, France), for at least 2 hours. The streptavidin was washed and the slice incubated in 2 mg/ml diaminobenzidine in Tris buffer, to which 0.002% H_2_O_2_ was added to initiate the staining reaction. Nickel intensification was occasionally employed (0.15% nickel ammonium sulphate). The reaction was stopped by washing in PBS. The slice was then mounted in Mowiol and, once it had set, the slice was observed. The pictures shown in Fig. 2 were taken using a 20x dry objective and a 63x oil-immersion objective.

**Fig. 1.**
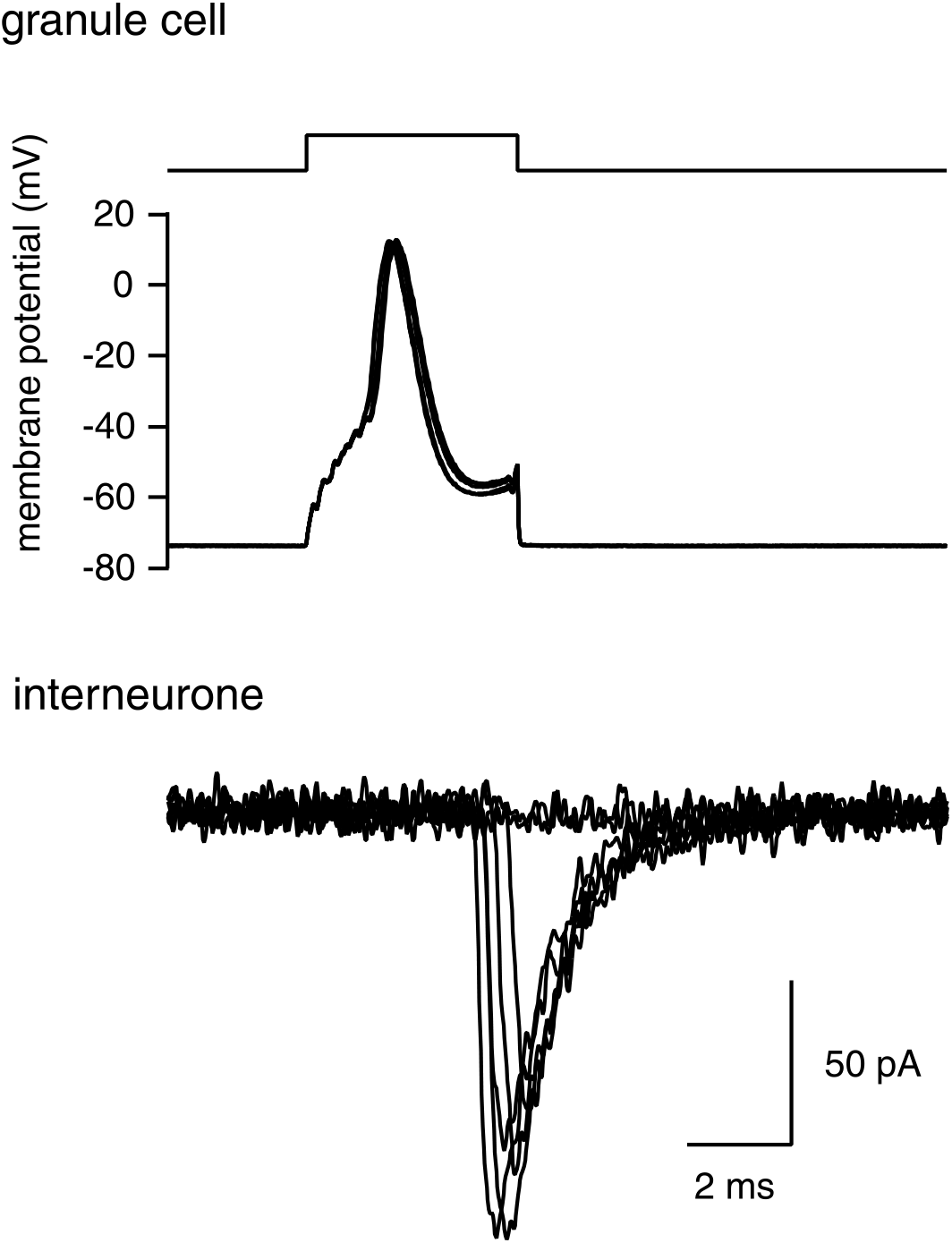
Specimen recordings from synaptically connected granule cell and interneurone. The top panel shows the granule cell’s membrane potential, which was clamped at -73 mV, except for a period of about 4 ms, when the granule cell’s amplifier was switched to current clamp and a depolarising current was injected (timing indicated by the step function at top) to elicit an action potential. In the lower panel, the corresponding postsynaptic responses (EPSCs) recorded under voltage clamp in the interneurone are shown.

**Fig. 2.**
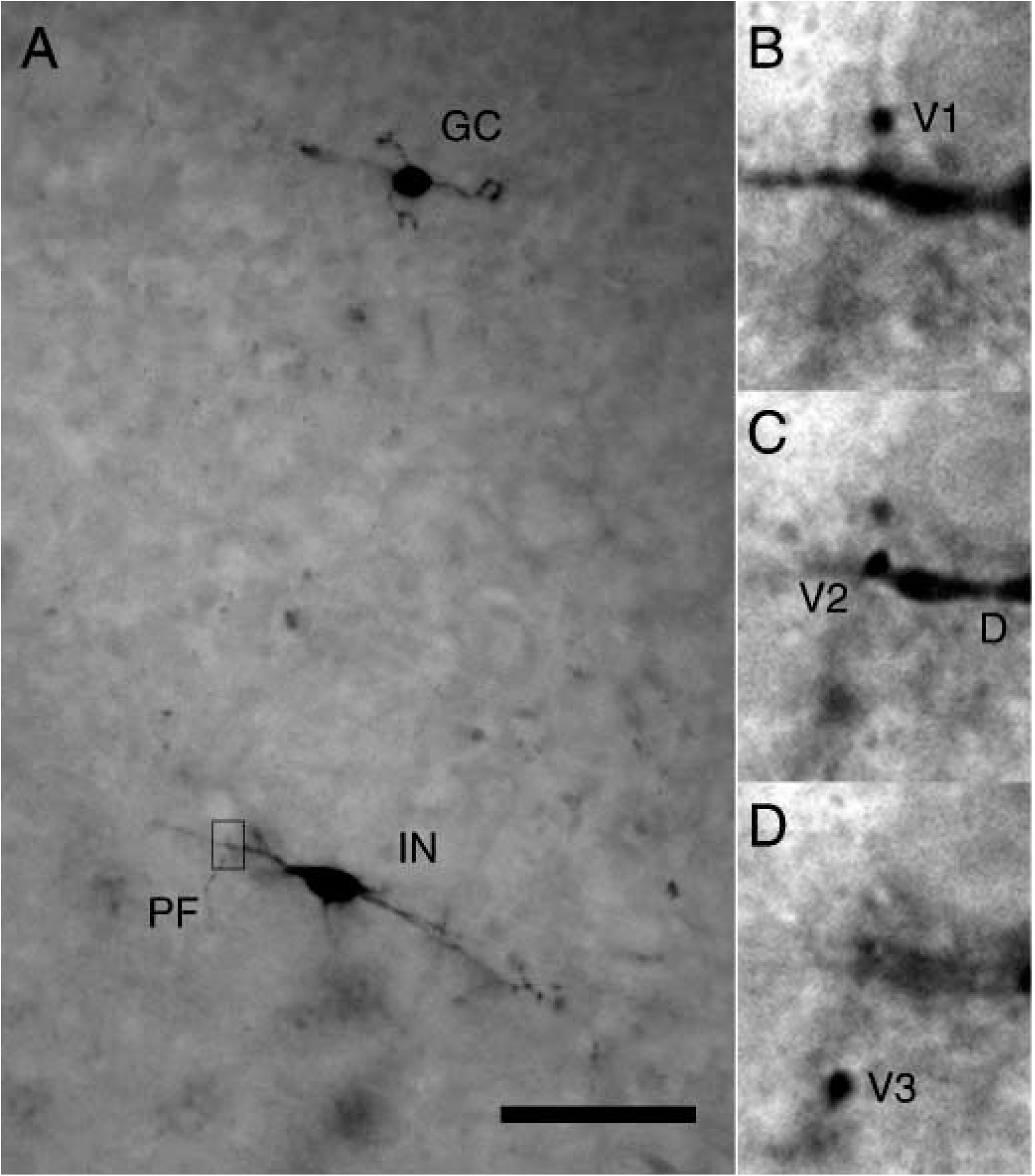
Morphology of a granule cell-interneurone synaptic connection. The biocytin labelling of the pair of Fig. 1 was successfully developed and allowed identification of a single point of contact between the two cells. **A**. Broadfield montage showing both the granule cell (‘GC’) and the interneuron (‘IN’). The relative positions of the cells have been preserved, but they were photographed on slightly different focal planes. The parallel fibre (‘PF’) is barely visible at this magnification as it crosses the plane of the interneurone dendrites. Three higher magnification views of the contact region (upright rectangle) at different focal planes are shown in B, C and D. **B**. The parallel fibre varicosity (‘V1’) preceding (nearer the granule cell soma) that making the recorded connection. **C**. The varicosity (‘V2’) making the synaptic contact rests on the interneurone dendrite (‘D’), which runs orthogonal to the parallel fibre. **D**. The varicosity (V3) succeeding the contact. Scale bar: 50 µm for A and 6 µm for B, C and D.

### Automatic detection of EPSCs and analysis

We used an automatic method for EPSC detection in order to make this process objective and reproducible. The particular method used was chosen under the constraint that it should perform reasonably in detecting EPSCs superposed upon the decay phase of others. We developed a simple method based upon the second derivative of the current trace. This detects changes of slope and is therefore fairly insensitive to the pre-existing current slope. The putative quantal events we wished to detect displayed the following features: they were monophasic inward currents; the ‘rising’ phase lasted a few hundred micros; the decay phase was roughly exponential, with a time constant of the order of 1ms. Because derivatives are noisy, we filtered the current trace first (box filter of 7--11 data points). Then the following quantity was calculated (which resembles the central difference approximation of the second derivative):

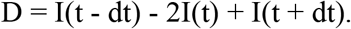

dt was adapted to the the rise-time of the EPSCs, the key EPSC feature exploited by the detection algorithm. In general, a dt of around 400 µs was used. Finally, it was noted that EPSCs thus transformed generated characteristic biphasic waveforms whose minima and maxima reflected the slope changes at the beginning and the end of the rising phase, respectively. This last feature was exploited in a second detection stage that improved the sensitivity of EPSC detection somewhat. A difference was formed between points separated by dt on D. This formed the measure used to detect EPSCs. When the measure crossed a threshold (20 to 30pA) in the positive-going direction, an EPSC was detected. It was timed as the following zero crossing of D (this corresponds to a point near the middle of the rising phase of the EPSC). Once the parameters had been adjusted by inspection for a particular cell, all EPSCs in that cell were detected using the same settings. The settings were conservative, to eliminate false positives.

The comparison between two Poisson rates was performed using the on-line Java calculator provided by SGCORP (http://www.sgcorp.com).

## Results

Specimen traces from a synaptically connected granule cell-interneurone pair are shown in Fig. 1. The upper trace shows the granule cell’s membrane potential, which was clamped at -73 mV between stimuli. Stimulation was effected by switching to current-clamp and injecting a depolarising current. The stimulating current was adjusted so that a single action potential was reliably elicited, as shown. The lower traces show the corresponding excitatory postsynaptic currents (EPSCs) recorded in the voltage-clamped interneurone. These EPSCs were mediated exclusively by AMPA receptors, since the recordings were carried out in the presence of 1 mM Mg ions and 50 µM D-APV. Their rapid rise, less than 200 µs (20--80%) in the cell illustrated, for simple events like those shown, suggests that the time resolution of the recordings is more than satisfactory for a dendritic input. The decay time constant (about 1 ms) is very close to that measured for deactivation of responses in patches excised from the same cells (Barbour et al., 1994).

The recorded pairs were routinely filled with biocytin. The revelations of the recorded cells attempted after the experiments were in the main unsuccessful, presumably because one or the other cell was often damaged at (or before) the end of the recording, leading to dispersal of the biocytin and consequent faint or absent staining, as well as giving dendrites a varicose appearance. Nevertheless, in the three successfully filled pairs, what appeared to be a single axo-dendritic synaptic contact between a parallel fibre varicosity and an interneurone dendrite was indentified; one of them is shown in Fig. 2.

Observation of the EPSCs evoked in the interneurones by single action potentials in the granule cell revealed quite frequent occurrences of compound EPSCs that clearly resulted from the superposition of two or more quantal EPSCs with slightly differing times of arrival. Some examples are shown in Fig. 3. Because these synaptic connections are very likely to involve just a single contact (the examples shown are from the pair in Fig. 2), these multiple release events suggest that more than one vesicle can be released per action potential at these synapses. In order to confirm or disprove this hypothesis, we sought to test whether such compound release events could reflect the chance arrival of spontaneous EPSCs from other parallel fibre synapses on the same interneurone.

**Fig. 3.**
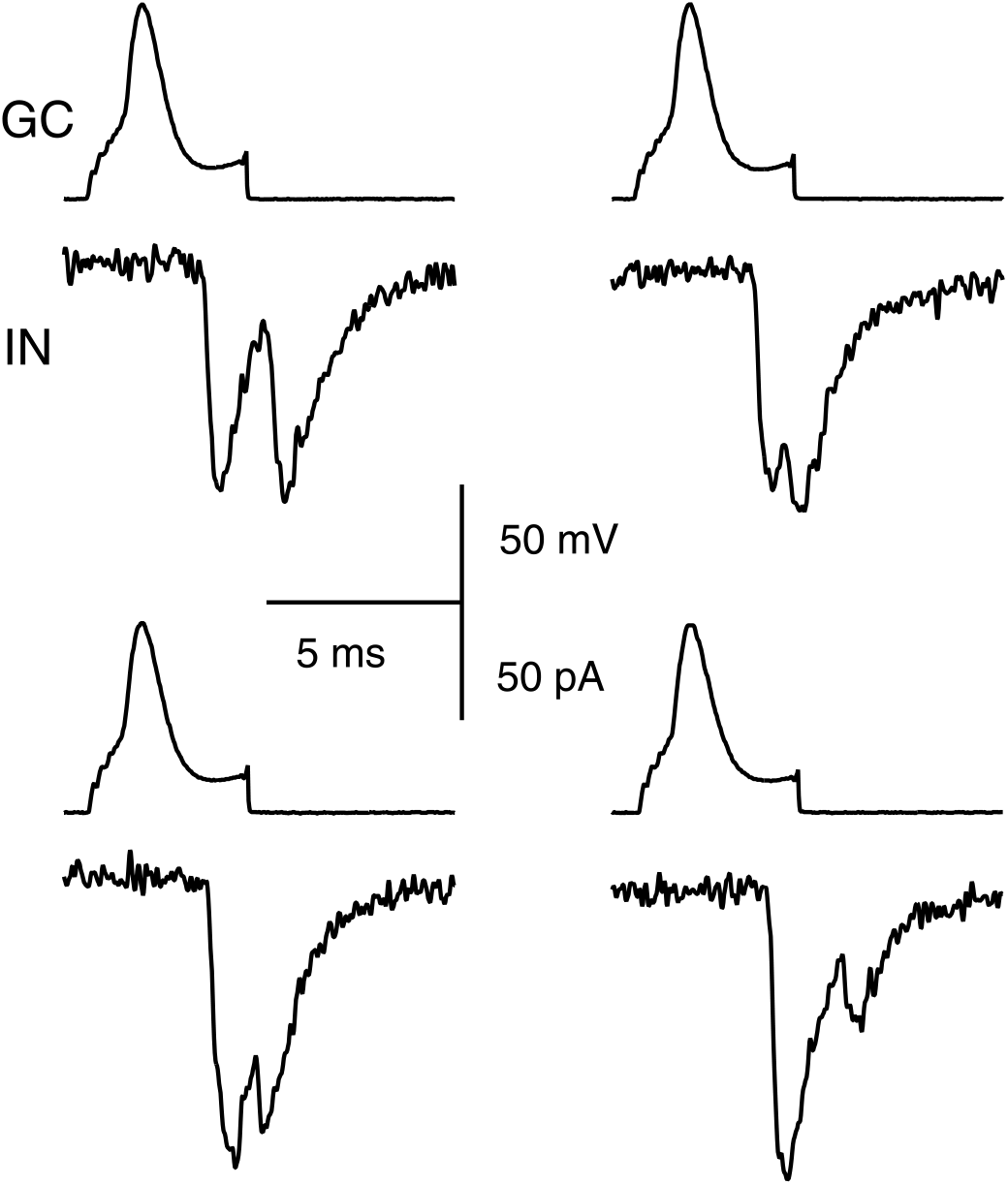
Multiple release events per action potential are frequently observed at granule cell-interneurone contacts. Examples of compound or multiple release events that appear to be triggered by the preceding action potential. In each of the four sweeps (all taken from the pair of Figs. 1 and 2), the upper trace shows the granule cell membrane potential and the elicited action potential, while the lower trace shows the EPSCs recorded in the interneurone.

Although in the example traces shown in Fig. 3, the presence of more than one release event seems very clear to the observer, a continuum of cases was observed. Numerous EPSCs appeared to result from the summation of nearly-synchronous simple events, but this was indicated by more subtle features of the EPSC, such as an inflection on the rising phase (see the bottom left EPSC in Fig. 3 for a very subtle example). Although the experienced eye of the experimenter is probably a sensitive detector of multiple release events, we wished to use a more objective, automatic method, to enable conclusive application of a statistical test. Some previously-used methods of automatic EPSC detection are not particularly well-suited to the particular task of detecting superposed EPSCs. Although detecting and fitting individual EPSCs using an EPSC-shaped template is probably the most accurate method, disentangling compound events (see Fig. 3A) automatically would be complex and was not attempted. Simpler methods generally involve thresholding a measure related to the first derivative of the trace (usually the difference between two short averages separated by an interval). However, this approach has a serious shortcoming if one wishes to detect EPSCs superposed upon the decay phases of preceding EPSCs. The decay of a preceding EPSC makes it more difficult for a second EPSC to attain the threshold ‘slope’. This would lead to an artefactual undercount of superposed EPSCs.

To avoid this problem, we developed a method based upon a measure related to the second derivative of the trace. The second derivative reflects the ‘rate of change of slope’ and should therefore be much less sensitive to superposition of quantal EPSCs. The algorithm is described in the Methods; its application to the corresponding EPSCs in Fig. 3B resulted in the detection of two release events in each case. The parameters of the detection algorithm were set conservatively, to eliminate false positives. More subtle indications of EPSC superposition did not result in the indentification of multiple EPSCs.

Using this EPSC detection algorithm, we counted spontaneous EPSCs that occurred in the period immediately before the first action potential of each sweep and those detected in a brief time window after each action potential. A window of 4 ms sufficed to enclose most of the evoked EPSCs. In 19/20 pairs recorded, the frequency of double detections following an action potential exceeded the frequency of spontaneous EPSCs. (The frequency is simply the number of EPSCs detected divided by the total time of the windows analysed). The average frequency of spontaneous EPSCs was 0.44 ± 0.75 Hz (n = 19 pairs), while the mean frequency of double EPSCs within a 4 ms window following the action potential was 20 ± 18 Hz. Total times of 2.4 -- 15.7 s (spontaneous EPSCs) and 0.60 -- 2.1 s (evoked EPSCs) were analysed. In the remaining cell, the release probability was very low and no multiple events were observed.

We now test to what extent this difference can be considered as evidence that the multiple release events are triggered by the action potential or whether they could reflect random spontaneous EPSCs. For each synaptic connection we tested the null hypothesis that no more than a single quantal could be triggered by an action potential, and that spontaneous EPSCs occur at random, according to a Poisson process. In this situation, double (or triple etc) EPSCs must arise from the chance superposition of a single released EPSC from the synapse being stimulated and spontaneous EPSCs from other synapses. The test must decide what is the probability that a single Poisson process could give rise to the different frequencies measured for spontaneous EPSCs during the control period and multiple EPSCs following the action potential. The null hypothesis is thus that the underlying Poisson parameters (rates or frequencies) are equal for the two periods. The alternative hypothesis is that the two samples are drawn from populations with different Poisson rates (this is the two-tailed test).

As an example, we take the data for the connection shown in Figs. 1, 2 and 3. During control periods before the first action potential of each sweep, totalling 6.303 s, 11 spontaneous EPSCs were detected. During 4 ms windows beginning 1 ms after each action potential (this latency was adjusted for each pair), 11 double and 1 triple release events were detected in 1.472s. The EPSC frequencies to be compared are then 1.7Hz for the control period and 8.8 Hz after the action potential. Application of a two-tailed test for comparing two Poisson rates leads to the rejection of the null hypothesis (P< 0.0001). In the two other successfully labelled pairs, similar conclusions were reached (P = 0.0007, P < 0.0001). Indeed, in 19/20 of the pairs recorded, the null hypothesis was rejected (range P < 0.035 to P < 0.0001). We therefore conclude that multiple release events can occur following a single action potential at the synaptic connections between granule cells and interneurones.

## Discussion

We have shown that multiple release events often occur in rapid succession following a single action potential at parallel fibre-molecular layer interneurones in immature rat cerebellar slices at room temperature. This conclusion was individually confirmed for 19/20 of the pairs recorded, including three for which a single synaptic contact point had been identified.

The presynaptic varicosity on the parallel fibre may contact the dendritic shaft or spines of stellate cells and basket cell (Palay and Chan-Palay, 1974). Somatic contacts are rare but do occur. The orthogonal arrangement of the presynaptic fibre and the planar postsynaptic dendrites (Introduction), coupled with the morphology of the small number of our cell pairs that were successfully labelled, strongly suggest that most of these synaptic connections consist of a single contact. However, electron microscopic studies of parallel fibre-interneurone synapses in the adult mouse (Lemkey-Johnston and Larramendi, 1968) show (their Fig. 17/Plate 9) that varicosities can make two synapses simultaneously with a spine and a dendrite from the same interneurone. It is possible that we would have failed to detect such a situation in our filled pairs. The frequency of such double synapses is unknown, although the work of Napper and Harvey (Harvey and Napper, 1988; Napper and Harvey, 1988a, b; Harvey and Napper, 1991) has shown that 80% of parallel fibre varicosities only make a single synaptic contact; many of the varicosities that make two contacts do so with two Purkinje cell spines. Therefore, although we cannot rule out the possibility that some of our recordings involved double synaptic contacts, we consider it unlikely to be the case for all of our recordings.

Subject to the above caveat, we conclude that the multiple release events we observe are intrinsic to the operation of a single contact. The obvious and most likely explanation for these multiple release events is that each corresponds to the release of a vesicle of glutamate. In this case release at this synapse is ‘multivesicular’. However, we do not completely exclude the possibility that a single vesicle might release its contents in several stages. Transmitter release is thought to begin with the opening of a protein channel between the vesicle lumen and the synaptic cleft. In large vesicles, this ‘fusion pore’ can close without full fusion occurring (Albillos et al., 1997). Such detailed information is not available regarding the small synaptic vesicles that release glutamate, but analogous behaviour could possibly generate multiple release events from a single vesicle.

Modifications of synaptic strength are considered to play a fundamental role in most learning and memory processes. Our results have some bearing on possible ‘presynaptic’ mechanisms of synapse modification. If only a single vesicle could be released per action potential, presynaptic mechanisms could only change the probability of its release. This would have the important consequence that precise read-out of the state of the synapse would require an averaging stage. If, however, we assume that one vesicle does not saturate the postsynaptic receptors [(Silver et al., 1996; Forti et al., 1997; Liu et al., 1999; McAllister and Stevens, 2000; Barbour, 2001); it also does not appear to be the case at the present synapse], then multivesicular release could allow a finer read-out of synaptic strength for a single action potential. Ultimately, this would increase the amount of information that could be stored in a synapse.

## Acknowledgements

We thank Philippe Ascher and the members of the Laboratoire de Neurobiologie for their generous support.

